# Ca^2+^-dependent liquid-liquid phase separation underlies intracellular Ca^2+^ stores

**DOI:** 10.1101/2021.07.06.451223

**Authors:** Joshua E. Mayfield, Adam J. Pollak, Carolyn A. Worby, Joy C. Xu, Vasudha Tandon, Mainak Bose, Allison McNeilly, Sourav Banerjee, Alexandra C. Newton

## Abstract

Endoplasmic/sarcoplasmic reticulum Ca^2+^ stores are essential to myriad cellular processes, however, the structure of these stores is largely unknown and existing models do not address all literature observations. We investigate CASQ1 - the major Ca^2+^ binding protein of skeletal muscle - and discover Ca^2+^-dependent liquid-liquid phase separation activity. The intrinsic disorder of CASQ1 underlies this activity and is regulated via phosphorylation by the secretory pathway kinase FAM20C. This divalent cation driven condensation demonstrates liquid-liquid phase separation occurs within the endoplasmic/sarcoplasmic reticulum, mechanistically explains efficient Ca^2+^ buffering and storage, and represents a largely unexplored mechanism of divalent-cation driven protein association.

## Main Text

Compartmentalization of molecules and metabolic processes is essential to life. In eukaryotes, membrane-bound organelles are well-studied and facilitate cellular functions including the storage and replication of genetic material, establishment of concentration gradients, and sequestration of cytotoxic entities and chemistries (Aguzzi and Altmeyer, 2016; Alberti et al., 2019; Dutagaci et al., 2021). In addition to membrane-bound organelles, membrane-less organelles (MLOs) are observed in the nucleus and cytoplasm and include the nucleolus, nuclear speckles, stress granules, and others (Aguzzi and Altmeyer, 2016; Dutagaci et al., 2021; Feric et al., 2016; Galganski et al., 2017; Protter and Parker, 2016). These biological condensates form via liquid-liquid phase separation (LLPS) wherein biomolecules enter a dense liquid-like phase distinct from the surrounding aqueous environment. These liquid-like droplets act to increase the local concentration of biomolecules, potentiate metabolic outcomes, and sequester components that would otherwise freely diffuse. Both proteins and RNA are known to undergo LLPS *in vivo*. Protein LLPS relies upon amino acid composition, conformational flexibility, and environmental conditions including temperature, pH, ionic strength, and the presence of other macromolecules like RNA (Aguzzi and Altmeyer, 2016; Alberti et al., 2019; Dutagaci et al., 2021; Wang et al., 2019). Observed mechanisms of RNA LLPS rely upon the presence of polycations, generally proteins, that can interact with polyanionic RNA and induce LLPS via coacervation – a specific form of interaction between ionic polymers (Alberti et al., 2019; Dutagaci et al., 2021; Sing, 2017). Less is known concerning the ability of polyanionic proteins to undergo LLPS through similar charge-dependent mechanisms. These proteins are particularly abundant in the nucleus, where MLOs are well established, but also in the endoplasmic/sarcoplasmic reticulum (ER/SR) where their negative charge is thought to facilitate high-capacity ionic calcium (Ca^2+^) storage (Supplementary Figure 1) (MacLennan and Reithmeier, 1998; Royer and Ríos, 2009). However, the ability of these proteins to undergo LLPS remains unexplored.

The Ca^2+^-sequestering protein, Calsequestrin-1 (CASQ1) is a polyanionic protein that has features allowing it to serve as a model protein for addressing LLPS. CASQ1 is the most abundant Ca^2+^ binding protein of the SR of skeletal muscle where it binds the majority of lumenal Ca^2+^. CASQ1 is observed in the ER of other tissues including cerebellar Purkinje neurons (2013; Woo et al., 2020). CASQ1 is highly conserved amongst vertebrates, binds calcium with high capacity (50-80 Ca^2+^/molecule) and low affinity (*K_d_* ∼ 10^3^ M^-1^), and is characterized by low iso-electric point and an abundance of acidic residues (Sanchez et al., 2012; Woo et al., 2020). Multivalent Ca^2+^ binding induces reversible oligomerization to drive total ER/SR Ca^2+^ stores well into the millimolar range, while buffering free lumenal Ca^2+^ concentrations to consistent low millimolar levels (Woo et al., 2020). Mutation of CASQ1 results in human myopathies and deletion in animal models results in defects in skeletal muscle function and Ca^2+^ handling (Barone et al., 2017; Lewis et al., 2015; Olojo et al., 2011; Paolini et al., 2007; Woo et al., 2020). This reversible oligomerization is central to excitation-contraction coupling in skeletal muscle as the oligomeric and non-oligomeric forms of CASQ1 differentially regulate Ca^2+^ release from the SR through physical association with ryanodine receptors (RyR) (Manno et al., 2017; Royer and Ríos, 2009; Woo et al., 2020). Therefore, a great deal of interest exists surrounding the physical structure of oligomeric CASQ1 as the assemblage underlies both lumenal Ca^2+^ stores and regulates Ca^2+^ release.

X-ray crystallography suggests CASQ1 is highly structured and oligomerizes into linear polymers (MacLennan and Reithmeier, 1998; Sanchez et al., 2012; Wang et al., 1998). However, this linear polymer model does not explain structures observed *ex vivo* and in cell culture. Electron microscopy (EM) of fixed and dehydrated muscle tissue from northern water snakes demonstrate CASQ1 forms a gel-like matrix *in situ*, characterized by repeated nodal points (Perni et al., 2013). Electron tomography of frozen-hydrated triad junctions reveals a variety of oligomerized states including spherical Ca^2+^-rich bodies (Wagenknecht et al., 2002). Immunolabeling of differentiated primary myotubes derived from mice reveal endogenous CASQ1 forms distinct spherical puncta like structures (Felder et al., 2002). EM studies of non-muscle tissues, specifically Purkinje neurons, reveal non-linear CASQ1/Ca^2+^ rich-structures known as calciosomes (Takei et al., 1992; Villa et al., 1991; Volpe et al., 1993). In cell culture, immunolabelling and EM of stably transfected L6 myoblasts reveal CASQ1 and Ca^2+^ form dense spherical vacuoles while live cell imaging of HeLa cells overexpressing fluorescent protein fusions of CASQ1 yield a variety of oligomeric forms including linear aggregates and spherical bodies (Barone et al., 2017; Gatti et al., 1997). Finally, in solution structural approaches under physiological conditions suggests monomeric CASQ1 in the Ca^2+^-free state is highly expanded (Cozens and Reithmeier, 1984). This contrasts with the compacted structures observed in crystals demonstrating even this fundamental unit is poorly understood. In context, the crystallographic data do not account for the diverse structures observed under physiological conditions. We surmise the discrepancy between crystallographic and physiological data is the result of the non-physiological conditions under which CASQ1 crystallizes.

Here we demonstrate CASQ1 oligomerizes as LLPS bodies. Utilizing negative stain EM, live-cell fluorescent microscopy, and time-resolved brightfield microscopy, we directly observe CASQ1 LLPS under physiological conditions. Through application of in solution structural methods we demonstrate the intrinsic disorder of CASQ1 influences its LLPS capacity by potentiating entry into the LLPS state. We show this intrinsic disorder is regulated by the secretory pathway kinase FAM20C, which phosphorylates structurally conserved regions of CASQ1 to increase disorder. Finally, we develop a homo-FRET based genetically encoded biosensor to monitor CASQ1 oligomerization in live cells. Collectively, our data reveal a novel mechanism of polycationic protein LLPS, establish LLPS occurs within the ER/SR, and point to a potentially widespread activity of intrinsically disordered acidic proteins found throughout eukaryotic proteomes. These data explain mechanisms underlying ER/SR Ca^2+^ storage and release and account for both biochemical and physiological data surrounding CASQ1.

## Results

### CASQ1 oligomerizes via LLPS

To address discrepancies between CASQ1 structures observed *in vitro* using crystallography and in *ex vivo* and cellular environments we first performed negative stain EM under lower ionic strength conditions and at physiological pH for purified CASQ1. CASQ1 was oligomerized through the addition of CaCl_2_, plated onto EM grids, and negatively stained with uranyl acetate. Resultant EM micrographs revealed densely negatively stained structures on the nanometer scale (Figure 1A). These structures contained spherical globules interconnected by thinner stretches of density and were qualitatively similar to the nodal structures described in both fixed muscle tissue and frozen hydrated triad junctions (Perni et al., 2013; Wagenknecht et al., 2002). The structures observed were larger than those seen in fixed tissue, potentially due to the dehydrating cryoprotection and prolonged incubations with highly ionic heavy metal stains required for EM of muscle tissue at such high resolution (Perni et al., 2013). In all cases, these structures were visually dissimilar to known linear protein polymer filaments (i.e. microtubules) (Johnson and Borisy, 1977). Interestingly, these oligomers contained branch points demonstrating that the nodal gel-like network observed *ex vivo* is an inherent property of CASQ1 and does not necessitate binding partners or other cellular factors (Perni et al., 2013).

**Figure 1.**
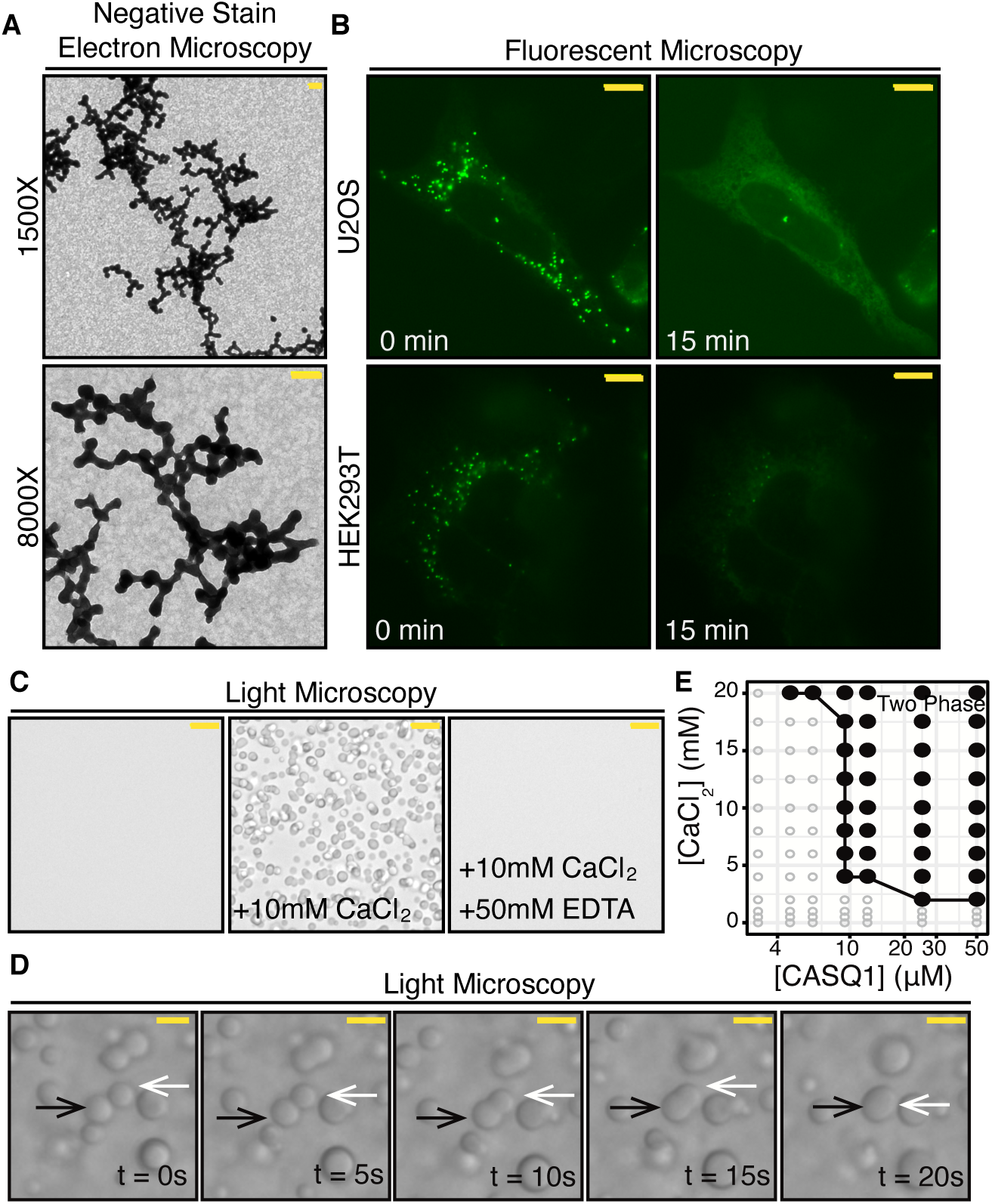
CASQ1 oligomerizes via LLPS. **(A)** Negative stain electron micrographs of CASQ1 oligomerized at 10mM calcium chloride and stained with uranyl acetate. (scale bar (yellow) = 200nm, magnifications indicated: 1500X (top), 8000X (bottom)) **(B)** Live cell fluorescent micrographs of U2OS (top) and HEK293T (bottom) cells at 0 (left) and 15 (right) minutes following thapsigargin treatment. (scale bar (yellow) = 10 μm). **(C)** Bright field micrographs of CASQ1 solutions prior to Ca^2+^ addition (left), following calcium chloride addition (center), and following EDTA addition (right). (scale bar (yellow) = 10 μm). **(E)** Phase diagram for CASQ1. Black filled circles indicate conditions under which CASQ1 undergoes liquid-liquid phase separation as indicated by the appearance of two phases. Black line indicates phase boundary. **(D)** Time resolved bright field microscopy of CASQ1 LLPS droplet fusion. The black and white arrow indicate droplets undergoing fusion (scale bar (yellow) = 2 μm (approximation)).

CASQ1 oligomerization has been visualized in cell culture, but observed structures are inconsistent. In stably transfected L6 myoblasts, HA-tagged CASQ1 generated dense spherical vacuoles where CASQ1 and Ca^2+^ were specifically accumulated (Gatti et al., 1997). Conversely, transient transfection of GFP-tagged CASQ1 into HeLa cells generated various structures including linear aggregates and rounded bodies when imaged in live cells (Barone et al., 2017). We replicated the GFP-tagged CASQ1 experiments with live-cell fluorescent microscopy in U2OS and HEK293T cells. U2OS and HEK293T cells were chosen as they express proteins relevant to SR Ca^2+^ handling (i.e. RyR1 and SERCA, Human Protein Atlas available from http://proteinatlas.org) (Uhlén et al., 2015). Additionally, HEK293 cells and their derivatives are used to study endogenous and overexpressed CASQ1 oligomerization dynamics (Wang et al., 2015). Overexpression of CASQ1-GFP yielded highly similar spherical puncta throughout the cell body of both cell lines (Figure 1B, U2OS (top-left) & HEK293T (bottom-left), 0 min). To confirm these structures were Ca^2+^-dependent oligomers and were properly localized to the ER/SR, we pharmacologically depleted Ca^2+^ from the ER lumen using thapsigargin, a specific inhibitor of sarco/endoplasmic reticulum Ca^2+^-ATPase (SERCA) (Sehgal et al., 2017). Upon treatment with thapsigargin there was a gradual dissolution of these puncta accompanied by a diffusion of the fluorescent species into a lacey intracellular network characteristic of ER (Figure 1B, U2OS (top-right) & HEK293T (bottom-right), 15 min). These data confirm that the assemblages observed in cell lines are not only reversible but also Ca^2+^-dependent and localized to the physiologically relevant organelle. Importantly, the spherical punctate structures observed in our live cell experiments were highly similar to those visualized at endogenous levels in differentiated primary myoblasts (Felder et al., 2002).

EM and live cell imaging both demonstrate CASQ1 oligomers form structures that are not explained by the linear polymer model. To directly visualize CASQ1 oligomerization in solution and circumvent any artifacts introduced by dehydration during EM we employed brightfield microscopy (BFM) of purified CASQ1 protein at physiological ionic strength (150mM) and pH (Figure 1C). In the Ca^2+^-free state solutions of CASQ1 were clear and had no distinguishing characteristics (Figure 1C, left). However, if oligomerization was induced through the addition of CaCl_2_, droplet-like structures readily appeared (Figure 1C, middle). These droplets were mobile, long lived (>1h), and displayed highly liquid-like characteristics. Time-resolved BFM demonstrated the rapid miscibility of these liquid-like droplets (Figure 1D), a hallmark of LLPS (Alberti et al., 2019). These droplets were completely dissolved upon the addition of Ca^2+^ chelator EDTA confirming they were both reversible and dependent upon free Ca^2+^ (Figure 1C, right). These bodies displayed additional characteristics of LLPS, including wetting, a process that occurs when the dense droplets sink and condense on the slide (Supplementary Figure 2) (Alberti et al., 2019). These condensed droplets maintain their miscible properties, observed as gradual growth and fusion over time (Supplementary Figure 2A to 2B). Collectively, these data suggest that oligomeric CASQ1 is a LLPS species at physiological ionic strength.

To confirm LLPS occurs across physiologically relevant concentrations of both protein and cation, we derived a phase diagram describing the dependence of LLPS activity on both Ca^2+^ and CASQ1 concentrations (Figure 1E). Total Ca^2+^ concentrations within the ER/SR have been quantified well into the millimolar range (21mM, (Fryer and Stephenson, 1996; Woo et al., 2020)). CASQ1 concentrations are high with some estimates placing it up to 100mg/mL (approximately 2mM) in the lumen of the SR (2013). LLPS activity was monitored using BFM at the Ca^2+^ and CASQ1 concentrations indicated and a distinct phase boundary - a clear distinction between concentrations at which CASQ1 does or does not undergo LLPS - was observed (Figure 1E). Even at modest CASQ1 concentrations (10μM) LLPS activity was readily observed for Ca^2+^ concentrations greater than approximately 3mM. Importantly, LLPS was not observed for Ca^2+^ concentrations less than 1mM. This approximates the physiological free Ca^2+^ concentration of the ER/SR and suggests these LLPS bodies efficiently buffer to this level. Taken together, the EM and live-cell imaging data show CASQ1 formed non-linear oligomerized bodies and BFM analysis of purified proteins demonstrated these arise from LLPS that occurs under physiologically feasible conditions.

### CASQ1 is an intrinsically disordered protein

A common characteristic of proteins that undergo LLPS is a degree of intrinsic disorder (Alberti et al., 2019). Crystal structures suggest CASQ1 is highly structured in both the Ca^2+^ free and bound states (Sanchez et al., 2012; Wang et al., 1998). However, in solution approaches demonstrate CASQ1 assumes an expanded structure (Cozens and Reithmeier, 1984). We hypothesized the condensed structures observed in crystals were the result of high ionic strength and/or low pH that neutralized CASQ1 surface charge to favor folding and crystallization. To test this, we performed a series of size exclusion chromatography (SEC) experiments of CASQ1 with varying concentrations of potassium chloride (KCl) to derive hydrodynamic properties including Stokes radii (Figure 2A & 2B). The Stokes radius of CASQ1 is highly dependent on the ionic strength of the buffer, ranged from 74Å at 0mM KCl to 40Å at 500mM KCl, and followed a logarithmic relationship (Figure 2B). To confirm this was not due to oligomerization of CASQ1 and was the result of structural expansion or contraction of the monomer we performed parallel analysis using tryptophan fluorescence spectroscopy. Tryptophan residues of any given protein readily fluorescence upon excitation in the UV range and the emission spectra varies depending on the polarity of the environment surrounding these residues. Tryptophan in a non-polar environment, for instance in the hydrophobic core of a protein or at a dimer interface, has a maximal emission wavelength of approximately 340-350nm. However, if tryptophan enters a polar environment, like the aqueous solvent, the emission spectra undergoes a red-shift and the maximal emission wavelength increases (Vivian and Callis, 2001). We excited solutions of CASQ1 at 295nm and monitored emission intensity from 320 to 400nm at varying KCl concentrations (Figure 2C). There was an obvious red-shift of the emission spectra as ionic strength decreased. Plotting the ratio of emission at 360/345nm clearly demonstrated the statistically significant relationship between emission ratio and KCl concentration (Figure 2D). Finally, CASQ1 was more susceptible to proteolytic degradation in low ionic strength conditions than in high ionic strength conditions (Supplementary Figure 3A). Collectively these data confirm that in solution CASQ1 is significantly structurally expanded compared to in crystal structures and this structural expansion is pronounced within physiologically relevant ionic strength ranges (100-200mM KCl, Figure 2D). To determine if the degree of structural expansion has functional consequences for CASQ1 LLPS, we performed turbidity assays monitoring CASQ1 oligomerization upon addition of CaCl_2_ to 10mM at various KCl concentrations. Turbidity assays are well-established in monitoring CASQ1 oligomerization and LLPS as the nanometer scale bodies scatter light at 350nm and can be monitored via absorbance at this wavelength (Alberti et al., 2019; Lewis et al., 2015). CASQ1 exhibited its highest oligomerization capacity at low KCl concentrations (0 & 50 mM, Figure 2E). As KCl concentrations increased (and the Stokes radii of CASQ1 decreased) there was a pronounced drop in absorbance at 350nm suggesting CASQ1 was less capable of undergoing LLPS in the structurally compacted state. The similar turbidity observed at 0 and 50mM KCl suggests the impact on turbidity was not fully explained by competition between potassium ions and Ca^2+^. Collectively, these data suggested structural expansion of CASQ1 influences its oligomerization capacity. Furthermore, it demonstrated that structural alterations in the Ca^2+^-free state inform Ca^2+^-dependent LLPS and represent a mechanism to regulate CASQ1 LLPS *in vivo*.

**Figure 2.**
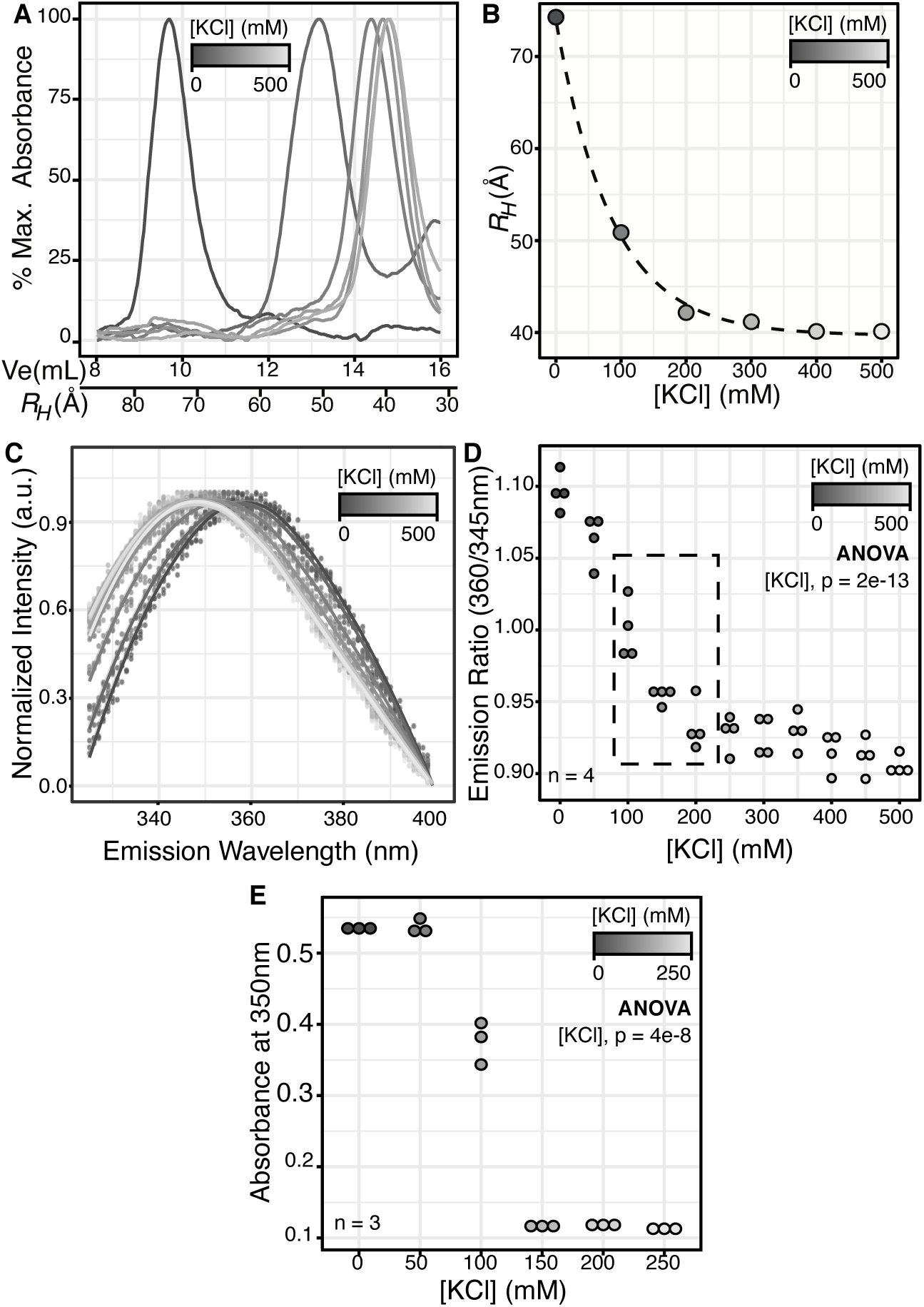
CASQ1 is an intrinsically disordered protein. **(A)** SEC elution profiles as monitored by absorbance at 280nm. Profiles obtained at 0, 100, 200, 300, 400, and 500mM KCl as indicated by greyscale. Data representative of three independent replicates. **(B)** Stokes radii (R_H_) determined from maximal peak intensity of Figure 2A. Stokes radii and KCl concentrations have a logarithmic decay relationship as indicated by fitted line. **(C)** Tryptophan fluorescence of spectra of CASQ1 (excitation = 295 nm, emission monitored 320-400 nm). KCl concentrations as indicated by greyscale. Points indicate individual data points; line indicates smoothed conditional mean. (n = 4). **(D)** Ratios of emission at 360 nm and 345 nm as a function of KCl concentration. Approximate physiologically ionic strength range (100-200mM KCl) indicated by dashed box. KCl concentration indicated by greyscale. Statistical significance of KCl on emission ration assessed using one-way ANOVA (n=4, One-Way ANOVA, F(1,42) = 112.8, p = 2e-13, alpha = 0.05) **(E)** CASQ1 turbidity assay as a function of KCl concentration as indicated by greyscale. Statistical significance of KCl concentration on turbidity assessed using one-way ANOVA (n=3, One-Way ANOVA, F(1,16) = 95.57, p = 4e-8, alpha = 0.05).

### FAM20C phosphorylates CASQ1 to increase intrinsic disorder and LLPS

CASQ1 is post-translationally modified via phosphorylation and glycosylation, but the modification enzymes responsible and the impact of these modifications on oligomerization are largely uncharacterized (Lewis et al., 2016). Our lab identified FAM20C as a ubiquitously expressed serine/threonine kinase localizing to the secretory pathway of eukaryotic cells, including the ER/SR lumen (Tagliabracci et al., 2012, 2015). We previously demonstrated the importance of FAM20C to muscle function in cardiac tissue where it phosphorylates a number of Ca^2+^ handling proteins (Pollak et al., 2018). We hypothesized CASQ1, which is expressed in both skeletal and cardiac muscle, is a substrate of FAM20C (Woo et al., 2020). To test this, we performed metabolic labelling of CASQ1 expressed alongside wild-type or catalytically inactive FAM20C in HEK293A cells grown in the presence of radioactive orthophosphate (^32^P) (Figure 3A), as previously (Pollak et al., 2018). These experiments demonstrated a robust incorporation of radioactive phosphate into CASQ1 when co-expressed with wild-type FAM20C. No radioactive incorporation was observed when CASQ1 was co-expressed with catalytically inactive FAM20C (D478A). Sequence analysis revealed two consensus FAM20C phosphorylation motifs (sequence = SXE, where X is any amino acid) at residues S248 and S369. Individual mutation of these residues to alanine reduced radioactive incorporation by roughly half and mutation of both essentially abolished ^32^P-phosphate incorporation (Figure 3A). *In vitro* phosphorylation of purified CASQ1 using purified FAM20C yielded identical results confirming this specificity (Supplementary Figure 4A). Residues S248 and S369 are highly conserved across mammals suggesting functional relevance (Figure 3B).

**Figure 3.**
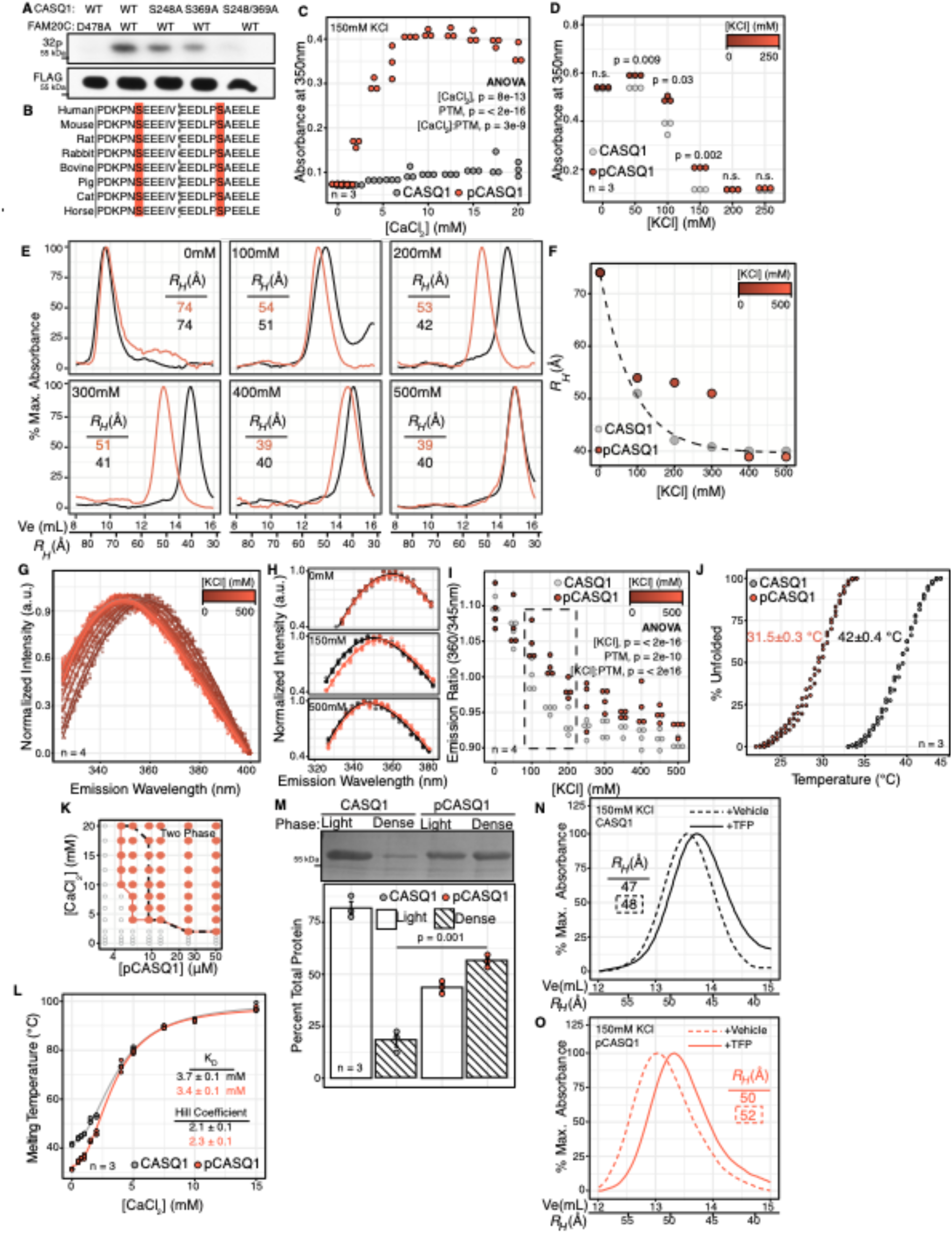
FAM20C phosphorylates CASQ1 to increase intrinsic disorder and LLPS. **(A)** Metabolic labeling of CASQ1 in the presence of radioactive orthophosphate (^32^P). **(B)** Amino acid sequence alignment of CASQ1 from various species. **(C)** Turbidity assay of CASQ1 (grey) and pCASQ1 (red) at 150mM KCl and various calcium chloride concentrations. Statistical significance of the influence of CaCl_2_, post-translational modification status (PTM), and their interaction on turbidity determined using two-way ANOVA (n=4, Two-Way ANOVA, ([CaCl_2_]: F(1,68)= 77.50, p = 8e-13, alpha = 0.05), (PTM: F(1,68) = 186.53, p = < 2e-16, alpha=0.05), ([CaCl_2_]:PTM: F(1,68) = 3e-9, alpha = 0.05) **(D)** pCASQ1 turbidity assay as a function of KCl concentration as indicated by dark to light red gradient. Unmodified CASQ1 data indicated in grey. Statistical significance determined by Welch’s t-test for CASQ1 and pCASQ1 data at a given KCl concentration (n=4, alpha = 0.05, n.s. = not significant). **(E)** SEC elution profiles as monitored by absorbance at 280nm. Profiles obtained at 0, 100, 200, 300, 400, and 500 mM KCl as indicated. CASQ1 traces indicated in black. pCASQ1 traces indicated in red. Stokes radii (*R_H_*) calculated from peak maxima indicated. Data representative of three independent replicates. **(F)** Stokes radii (*R_H_*) determined from maximal peak intensity of Figure 3E. For CASQ1 (grey) Stokes radii and KCl concentrations have a logarithmic decay relationship as indicated by fitted line. pCASQ1 (dark red to light red gradient) deviate from this relationship. **(G)** Tryptophan fluorescence of spectra of pCASQ1 (excitation = 295 nm, emission monitored 320-400 nm). KCl concentrations as indicated by dark red to light red gradient. Points represent independent readings; line indicates smoothed conditional mean. (n = 4). **(H)** Select tryptophan fluorescence spectra for CASQ1 (black) and pCASQ1 (red) at indicated KCl concentration. **(I)** Ratios of emission at 360 nm and 345 nm as a function of KCl concentration for pCASQ1 (dark red to light red gradient) and CASQ1 (grey). Approximate physiologically ionic strength range (100-200mM KCl) indicated by dashed box. KCl concentration indicated by dark red to light red gradient. Statistical significance of KCl concentration, post-translational modification status (PTM), and their interaction determined using two-way ANOVA (n=4, Two-Way ANOVA, ([KCl]: F(1,126) = 173.68, p = 2e-16, alpha = 0.05), (PTM: F(2,126) = 27.13, p = 2e-10, alpha = 0.05), ([KCl]:PTM: F(2,126) = 56.46, p = < 2e-16, alpha = 0.05). **(J)** DSF of CASQ1 (grey) and pCASQ1 (red) in the presence of SYPRO orange dye. Melting temperature indicated with error as standard error of the mean (n=4). **(K)** Phase diagram for pCASQ1. Red filled circles indicate conditions under which pCASQ1 undergoes liquid-liquid phase separation as indicated by the appearance of two phases. Red line indicates phase boundary for pCASQ1. Black dashed line indicates phase boundary for CASQ1 for reference. **(L)** CaCl_2_ dependence of CASQ1 (grey) and pCASQ1 (red) melting temperature. Data fit to modified Hill’s Equation to derive K_D_ and Hill Coefficient. CASQ1 values provided in black, pCASQ1 values in red. Error indicates standard error of fit values (n = 4). **(M)** Sedimentation assay of CASQ1 and pCASQ1. Coomassie stained SDS-PAGE gel of light and dense sedimentation fractions (top). Quantification of CASQ1 (grey) and pCASQ1 (red) percent in each sedimentation fraction (bottom). **(N)** SEC elution profiles for CASQ1 as monitored by absorbance at 280nm. Profiles obtained at 150mM KCl with or without 100μM TFP. Dashed line indicates vehicle control, solid line indicates TFP trial. Stokes radii (R_H_) calculated from peak maxima indicated. **(O)** SEC elution profiles for pCASQ1 as monitored by absorbance at 280nm. Profiles obtained at 150mM KCl with or without 100μM TFP. Dashed line indicates vehicle control, solid line indicates TFP trial. Stokes radii (R_H_) calculated from peak maxima indicated.

To determine the impact of FAM20C phosphorylation on CASQ1, we purified CASQ1 and phosphorylated it *in vitro* with FAM20C (pCASQ1). We first determined the impact of phosphorylation on Ca^2+^-dependent LLPS using a turbidity assay. At physiological ionic strength pCASQ1 displayed a significant increase in LLPS across CaCl_2_ concentrations greater than 2mM relative to CASQ1 (Figure 3C). Similarly, to CASQ1, pCASQ1 oligomerization was sensitive to ionic strength and was greater at low ionic strengths (0 and 50 mM) than at high ionic strengths (Figure 3D). Interestingly, pCASQ1 showed increased LLPS relative to CASQ1 at KCl concentrations from 50 to 150 mM but did not induce increased oligomerization at 0mM KCl (Figure 3D). We hypothesized these increases were due to structural expansion relative to CASQ1 and sought to test this using SEC (Figure 3E). We found that at KCl concentrations ranging from 100mM to 400mM, pCASQ1 had a larger stokes radius than CASQ1 and deviated significantly from the logarithmic relationship between Stokes radii and ionic strength observed for CASQ1 (Figure 3F). Importantly, at 0mM KCl CASQ1 and pCASQ1 had identical oligomerization capacity and displayed nearly identical SEC profiles (Figure 3E). To confirm that the increased Stokes radius was the result of structural expansion of the monomer and not phosphorylation-dependent oligomerization, we subjected pCASQ1 to tryptophan fluorescence spectroscopy (Figure 3G). The phosphorylated protein yielded similar results to CASQ1 with a clear red-shift in the emission spectra as ionic strength was decreased. Similar to the SEC experiments, tryptophan emission spectra at 0 and 500mM KCl were nearly identical, while spectra obtained at 150mM were different (Figure 3H). This red-shift was apparent in the emission ratios (360/345nm) where pCASQ1 also displayed the most pronounced structural expansion relative to CASQ1 at physiological ionic strength (150mM) (Figure 3I). This finding was further confirmed by increased susceptibility to proteolytic degradation of pCASQ1 relative to CASQ1 at identical ionic strengths (Supplementary Figure 3B).

To determine if this structural expansion is indicative of increased global intrinsic disorder, we performed differential scanning fluorimetry (DSF) in the presence of SYPRO-orange dye and determined CASQ1 and pCASQ1 melting temperatures (Figure 3J). Phosphorylation of CASQ1 resulted in a dramatic decrease in the melting temperature of the protein from 42°C to 31.5°C in the Ca^2+^-free state. This suggested phosphorylation of CASQ1 dramatically increased the disorder of this protein at physiological temperature. To investigate the importance of this structural expansion and intrinsic disorder to LLPS we derived a phase diagram similar to CASQ1 (Figure 3K). We found relative to the phase diagram for CASQ1 (Figure 1E), pCASQ1 had an extended phase boundary where it more readily underwent LLPS at lower protein concentrations. However, the minimal CaCl_2_ concentrations required were not altered suggesting pCASQ1 buffers to similar lumenal free Ca^2+^ levels. Therefore, we conclude FAM20C phosphorylation of CASQ1 induces structural expansion and confers disorder. This disorder potentiated entry into the LLPS state along physiologically relevant concentrations.

To mechanistically understand how phosphorylation influenced CASQ1 LLPS we first determined if pCASQ1 displays altered affinity for Ca^2+^. DSF is a sensitive assay for ligand binding as ligand bound proteins often increase in thermostability (Vivoli et al., 2014). Experiments were performed at 500mM KCl to minimize increases in melting temperature arising from oligomerization and isolate Ca^2+^ binding. Importantly, both CASQ1 and pCASQ1 melting temperatures similarly depended on Ca^2+^ concentration (Figure 3L). We fit these data to Hill’s Equation to approximate K_D_ (the ligand concentration at which half of the ligand binding sites are occupied) and Hill’s coefficient (indicative of cooperativity) and determined that these numbers were highly similar for CASQ1 and pCASQ1. These data suggested the increase LLPS is unlikely to arise from differences in Ca^2+^ affinity.

Given the roughly equivalent Ca^2+^ affinities of CASQ1 and pCASQ1, we hypothesized pCASQ1 more readily enters the LLPS state and investigated this using sedimentation assays. In sedimentation assays, LLPS is induced and the dense LLPS bodies are separated from the surrounding buffer via centrifugation. The dense phase is then resuspended in an equivalent volume to the light phase and protein is quantified (Wang et al., 2019). We found that under identical buffer conditions greater percentages of pCASQ1 entered the dense phase than CASQ1 (56% pCASQ1 vs. 18% CASQ1, Figure 3M). Collectively, these data demonstrated that, although CASQ1 and pCASQ1 may be similarly saturated with Ca^2+^, pCASQ1 more readily entered the LLPS state. This likely arises from its increased disorder and conformational flexibility - common characteristics of proteins that undergo LLPS (Alberti et al., 2019).

We were curious if known effectors of CASQ1 oligomerization exert their modulation via structural compaction/expansion. Anthracyclines are a common class of drugs that include the chemotherapeutic agents, Daunorubicin and Daunorubinicinol, as well as the antipsychotic, Trifluoperazine (TFP). Anthracycline therapy is associated with cardiac and muscular toxicity. Studies demonstrate these drugs bind CASQ1 to inhibit oligomerization and, ultimately, impact Ca^2+^ homeostasis (Charlier et al., 2005). Importantly, TFP preferentially binds CASQ1 in the Ca^2+^-free state and induces conformational change (Brown et al., 1994; He et al., 1993). TFP has little effect on Ca^2+^ saturated/oligomerized CASQ1 (He et al., 1993). This suggests TFP influenced oligomerization through modifications of the Ca^2+^-free conformation, similar to FAM20C-dependent phosphorylation. To monitor CASQ1 conformation in solution we applied SEC with or without TFP (Figure 3N & 3O). TFP induced structural compaction of both CASQ1 and pCASQ1 and supported structural compaction as a mechanism of CASQ1 oligomerization inhibition with clinical significance.

### Fam20C phosphorylation sites make unique contributions to CASQ1 disorder and LLPS

FAM20C phosphorylated CASQ1 at two sites, S248 and S369 (Figure 3A). To understand the contribution of each of these sites to increased intrinsic disorder and LLPS we generated and purified phospho-mimetic mutants S248E, S369E, and double mutant S248/369E. First, we monitored the behavior of these variants in a turbidity assay at physiological ionic strength with LLPS induced by 10mM calcium chloride (Figure 4A). Only S248/369E recapitulated pCASQ1 behavior, suggesting single phosphorylation events were not sufficient to increase LLPS. To determine if this is the result of impaired structural expansion, we monitored these mutants using SEC and found both S369E and S248/369E underwent expansion comparable to pCASQ1. S248E, on the other hand, had identical hydrodynamic properties to CASQ1 (Figure 4B). This suggested simple structural expansion was not sufficient to increase LLPS. We next monitored the impact of each site on melting temperature using DSF. We found that S248E and S248/369E displayed an identical melting temperature to pCASQ1 while S369E had an intermediate melting temperature between those of CASQ1 and pCASQ1 (Figure 4C). Neither structural expansion nor decreased melting temperature was sufficient alone to increase LLPS. CASQ1 must undergo both, via double phosphorylation, to gain appreciable LLPS activity. Therefore, both expanded hydrodynamic character and increased global instability are required.

**Figure 4.**
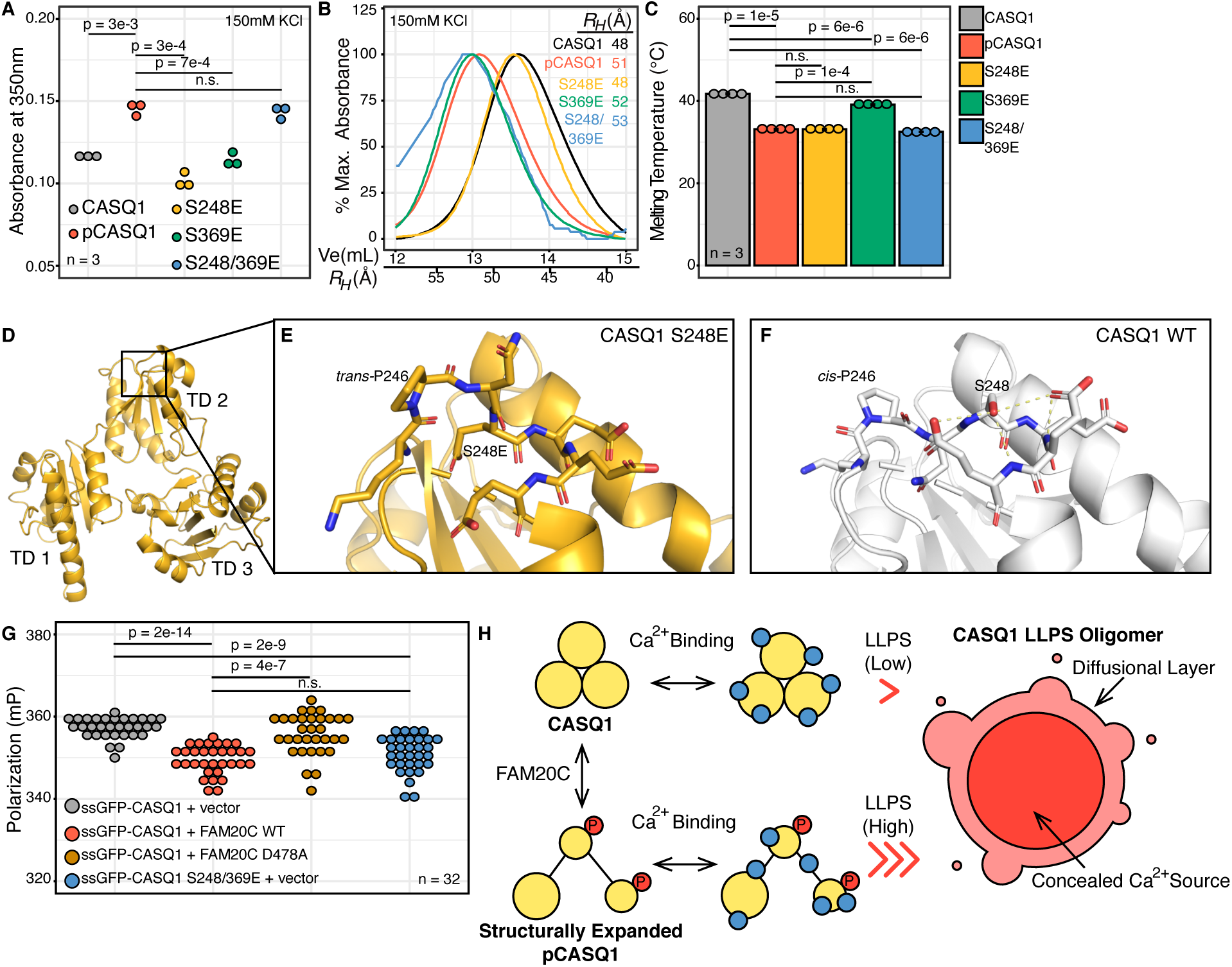
FAM20C phosphorylation destabilizes CASQ1 to influence CASQ1 LLPS *in vitro* and in cells. **(A)** Turbidity assay for CASQ1 (grey), pCASQ1 (red), S248E (yellow), S369E (green), and S248/369E (blue) at 10mM CaCl_2_ at 150mM KCl. Significance assessed using Welch’s t-test (n=3, alpha = 0.05, n.s = not significant). **(B)** SEC elution profiles for CASQ1 (grey), pCASQ1 (red), S248E (yellow), S369E (green), and S248/369E (blue) at 150mM KCl. Stokes radii (*R_H_*) calculated from peak maxima indicated. **(C)** DSF melting temperature assays for CASQ1 (grey), pCASQ1 (red), S248E (yellow), S369E (green), and S248/369E (blue). Statistical significance determined with Welch’s t-test (n = 4, alpha = 0.05, n.s. = not significant). Error bars indicate standard error of the mean. **(D)** Global view of S248E structure (yellow, PDB ID: 7MS4) with thioredoxin domains 1-3 indicated (TD1-TD3). **(E)** Local structural rearrangement proximal to the S248E site. P246 assumes a thermodynamically favorable *trans* conformation to stabilize the solvent exposed loop. **(F)** Local structure around S248 in wild-type structure (white, PDB ID: 5CRD). P246 assumes a thermodynamically disfavored *cis* conformation that is stabilized by hydrogen bonding (yellow) of S248 hydroxyl group. **(G)** Homo-FRET analysis of ssGFP-CASQ1 (grey), ssGFP-CASQ1 co-expressed with FAM20C WT (red), ssGFP-CASQ1 co-expressed with FAM20C D/A (orange), and ssGFP-CASQ1 S248/369E (blue). Statistical significance determined with Welch’s t-test (n = 32, alpha = 0.05, n.s. = not significant). **(H)** Model of FAM20C influence on CASQ1 LLPS. FAM20C phosphorylation does not impact Ca^2+^ binding but drives LLPS through increased disorder. The CASQ1 LLPS oligomer efficiently sequesters Ca^2+^ as only the surface of the body can freely diffuse into the aqueous lumen (labeled “Diffusional Layer”). Simultaneously, the Ca^2+^ sequestered to the center of the LLPS body is efficiently concealed (labeled “Concealed Calcium Source”).

### FAM20C phosphorylation destabilizes a conserved structural element in thioredoxin domains

To mechanistically explain the influence of phosphorylation on CASQ1 physical properties we investigated available crystal structures to understand what local characteristics were altered by phosphorylation or glutamate mutation. CASQ1 is composed of three thioredoxin domains, a common structural motif conserved from prokaryotes to mammals (Figure 4D). Structurally, they are characterized by a five beta sheet core surrounded by four alpha helices (Katti et al., 1990; Sanchez et al., 2012; Wang et al., 1998; Woo et al., 2020). S248 and S369 occur at structurally equivalent regions of thioredoxin domains two and three of CASQ1 (Supplementary Figure 5A-5D). These serines are located at the N-terminus of the fourth alpha helix of the thioredoxin domain and form a helix cap. Helix capping is a common structural feature where the peptide backbone is stabilized at the beginning or end of the alpha helix by hydrophobic or polar contacts (Aurora and Rose, 1998). Serine or other hydroxyl containing residues at this position are conserved from *E. coli* to humans, suggesting a significant structural contribution (Supplementary Figure 5E). In CASQ1, the hydroxyl groups of both S248 and S369 form a hydrogen bond with the amide of the peptide bond at the +3-residue position (Supplementary Figure 5C & 5D). These caps stabilize the loops extending from the fifth beta sheet and bolster the alpha helix, evidently stabilizing the protein and contributing to compaction. Based on the equivalently reduced thermostability, similar structural expansion, and identical oligomerization activity of pCASQ1 and S247/369E variant, we hypothesized that phosphorylation and glutamate mutation similarly disrupt these capping motifs and lend disorder to the protein. To determine local structural rearrangements, we performed x-ray crystallography experiments to visualize the phosphorylated or glutamate substituted sites. Despite extensive optimization, pCASQ1 yielded crystals characterized by poor diffraction and high mosacity that prevented determination of atomic resolution structures. Attempts to crystallize S369E and S247/369E were similarly complicated, likely owing to the entropy introduced by the structural expansion observed in SEC (Figure 4B). The S248E variant, however, yielded quality crystals that diffracted into the atomic resolution range despite its pronounced drop in melting temperature (Supplementary Table 1, PDB ID: 7MS4). Our crystallization conditions captured S248E as a dimer in the asymmetric unit. The individual monomers were highly similar to previously published structures (RMSD: Chain A: 1.0, Chain B: 0.8 relative to PDB ID 5CRD) (Wang et al., 1998). However, we observed substantial local rearrangement near the S248E mutation. Glutamate substitution disrupted helix capping and resulted in rearrangement of the loop connecting beta sheet five to alpha helix four of the second thioredoxin domain of CASQ1 (residues 244 – 247) (Figure 4E & 4F, Supplementary Figure 5F & 5G). This rearrangement appears to be driven by two structural features. First the side chain of glutamate is negatively charged and significantly larger than serine. This introduces electrostatic repulsion (from nearby E249, E250, and E251) and steric clashes. These factors shifted the mutated residue of S248E to a position occupied by N247 in wild-type structures atop a surface pocket formed by H191, A196, F197, and I229. The second key structural feature is the isomerization state of P246, a conserved residue in mammals (Figure 3B). In the wild-type structure, this proline assumed a thermodynamically disfavored *cis* conformation (Figure 4F) (Brandts et al., 1975). This conformation was stabilized by the hydrogen bonding network mediated by S248 at the helix cap. However, upon disruption of these interactions P246 converted to the more thermodynamically favored *trans* conformation (Figure 4E). This, coupled with the steric and electrostatic repulsion induced by S248E, displaced this loop by ∼7 Å into a solvent exposed position. The cis-to-trans isomerization of P246 acted as a thermodynamic spring to drive conformational change. This thermodynamically locked conformation likely contributed to the pronounced thermal destabilization observed for this mutant and highlights the importance of helix capping as a mechanism to stabilize thioredoxin domains.

### CASQ1 phosphorylation and intrinsic disorder influence LLPS in live cells

To understand the influence of FAM20C phosphorylation on CASQ1 oligomerization in live cells, we developed a method to quantify oligomerization in real time. Previous studies monitored secretory pathway oligomerization processes like insulin granule packing using homologous Förster Resonance Energy Transfer (homo-FRET) biosensors (Yi et al., 2015). These biosensors rely on FRET between identical fluorophores, and fluorophore proximity is indicated by decreased relative fluorescent intensity of light at incident polarization (Supplementary Figure 6A). Given the dramatic oligomerization observed for CASQ1-GFP (Figure 1B), we hypothesized similar constructs would be amenable to homo-FRET. We confirmed that GFP homo-FRET is an efficient indicator of fluorophore proximity in the secretory pathway by expressing the first 49 residues of CASQ1 (which contain the signal peptide) fused to GFP (ssGFP) and ssGFP fused to an additional GFP construct as a positive homo-FRET control (ssGFP-GFP) in HEK293A cells (Supplementary Figure 6B). Cells were harvested and the fluorescence polarization of equal numbers of cells were determined. There was a statistically significant decrease in polarization between ssGFP and the covalently linked ssGFP-GFP constructs, confirming that homo-FRET indicates GFP proximity within the secretory pathway (Supplementary Figure 6B). To determine if CASQ1 oligomerization can be monitored via homo-FRET in real time we repeated experiments performed in Figure 1B, with slight modifications. HEK293A cells were transfected with CASQ1-GFP, Ca^2+^ depleted through resuspension in Ca^2+^-free HBSS and transferred to an appropriate microplate. Cells were then treated with HBSS containing Ca^2+^ and either thapsigargin or vehicle control. Both groups display identical initial polarizations. Following treatment, a gradual drop in florescent polarization for the vehicle control was observed (Supplementary Figure 6C). Thapsigargin prevented this drop and maintained CASQ1 in a lower oligomerization state as indicated by nearly constant polarization levels (Supplementary Figure 6C). End point analysis of this same assay confirmed a statistically significant difference between the thapsigargin and vehicle treated cells (Supplementary Figure 6D). Importantly, lysis of the cells and chelation of the Ca^2+^ with EDTA normalized the homo-FRET signal confirming that differences were dependent on lumenal Ca^2+^ and CASQ1 oligomerization (Supplementary Figure 6E). This established homo-FRET as a means to monitor Ca^2+^-dependent CASQ1 oligomerization in cells.

To determine the impact of FAM20C on CASQ1 oligomerization we generated CASQ1 constructs containing an N-terminal GFP tag placed between CASQ1 residues 49 and 50 (ssGFP-CASQ1), similar to ssGFP constructs used in validation and for HA-tagged constructs used previously to study CASQ1 oligomerization (Wang et al., 2015). We chose this design to avoid potential artifacts resulting from tag proximity to the second and third thioredoxin domains where phosphorylation occurs and disrupts structure. Co-expression of ssGFP-CASQ1 with or without FAM20C in HEK293A cells revealed that FAM20C significantly decreased polarization at steady state Ca^2+^ levels (Figure 4G). Co-expression with a catalytically inactive FAM20C variant D478A did not result in significant decreased polarization and confirmed such decreases depended on FAM20C activity. To confirm that these drops in polarization result from modifications of CASQ1 and not from other substrates of FAM20C, we generated a ssGFP-CASQ1 S248/369E construct. Expression of this construct resulted in decreased polarization and was not statistically different from ssGFP-CASQ1 co-expressed with active FAM20C. Collectively, these data confirm the ability of FAM20C to increase CASQ1 LLPS under physiological conditions. The phospho-mimetic construct demonstrated that increased oligomerization results from alterations of CASQ1 not other FAM20C substrates. Collectively, these data support a model wherein FAM20C alters the intrinsic disorder of CASQ1 to influence its oligomerization through increased LLPS both *in vitro* and in cells (Figure 4H).

## Discussion

Data presented here provide a novel mechanism to regulate Ca^2+^ storage and release within the ER/SR via LLPS. We hypothesize this LLPS will be a widespread protein activity underlying numerous biological processes including Ca^2+^ signaling, neuronal function, neonatal nutrition, and biomineralization. Specifically, we demonstrated that CASQ1 is an intrinsically disordered protein with Ca^2+^-dependent LLPS activity under physiologically relevant conditions. This activity explains numerous biochemical and physiological observations made for CASQ1, which contributes to muscle function and various disease states. This activity allows CASQ1 to enter liquid-like droplets that explain the spherical bodies observed in live cells, hydrated triad junctions, and differentiated primary myoblasts (Barone et al., 2017; Felder et al., 2002; Wagenknecht et al., 2002). It can also account for the nodal gel matrix observed in fixed dehydrated muscle tissue (Perni et al., 2013). Although our data do not preclude the potential for linear polymers, these highly structured oligomers do not occur at physiological ionic strength. This conclusion is supported by multiple lines of evidence. First, CASQ1 crystal structures do not account for the conformations observed in solution under physiological conditions (Cozens and Reithmeier, 1984). Indeed, we only observed CASQ1 in a compacted state comparable to that observed in crystal structures at non-physiologically high ionic strengths. Similarly, we demonstrated factors that increase CASQ1 intrinsic disorder, like FAM20C phosphorylation, drive oligomerization *in vitro* and in live cells. Small molecules known to inhibit CASQ1 oligomerization *in vivo* appear to decrease its degree of intrinsic disorder in exerting their inhibitory effect (Charlier et al., 2005; He et al., 1993). These modulations of activity are counterintuitive to a linear polymer model necessitating highly structured CASQ1. Additionally, LLPS may contribute to the CASQ1 association/dissociation thought to occur in a contraction-to-contraction manner in muscle tissue (Manno et al., 2017; Woo et al., 2020). The linear polymer model necessitates the correct orientation of Ca^2+^ saturated CASQ1 with the growing oligomer. This represents a potentially rate limiting kinetic challenge, similar to other orientation-intensive processes like actin filament nucleation and growth (Firat-Karalar and Welch, 2011). LLPS circumvents this allowing for oligomerization regardless of orientation and is ideal for rapidly exchanged CASQ1 oligomers and the dynamic nature of Ca^2+^ signaling.

Alterations in CASQ1 intrinsic disorder may explain mutations linked to human disease. Disease mutations are known to influence CASQ1 oligomerization *in vitro* and *in vivo*, however these effects are not explained in the context of available crystal structures (Barone et al., 2017; Lewis et al., 2015; Sanchez et al., 2012). For instance, multiple mutations of CASQ1 decrease its oligomerization capacity and result in Myopathy, tubular aggregate, 1 (TAM1). These mutations include D44N, G103D, and I385T. Importantly, these mutations do not align with interaction interfaces observed in crystal structures suggesting alternative mechanisms are at play (Barone et al., 2017; Sanchez et al., 2012). We hypothesize that subtle alterations in CASQ1 intrinsic disorder and LLPS activity may inhibit oligomerization and drive TAM1. Additionally, a mutation proximal to S248, D244G, induces a disease state known as Myopathy, vacuolar, with CASQ1 aggregates (VMCQA) and dramatically increases oligomerization *in vitro* (Lewis et al., 2015). We hypothesize inappropriate introduction of the flexible amino acid glycine proximal to the helix cap increases the intrinsic disorder of this region and circumvents regulatory mechanisms described here to drive aberrant oligomerization. Therefore, mutations influencing the intrinsic disorder and LLPS of CASQ1 represent an unexplored paradigm for understanding associated myopathies in biochemical terms.

LLPS is an attractive model to explain ER/SR Ca^2+^ storage. Previous publications demonstrate that RyRs access a concealed, reversible, and high-capacity Ca^2+^ source that does not rapidly equilibrate with the aqueous lumenal environment (Guerrero-Hernandez et al., 2010; Perez-Rosas et al., 2015; Sánchez-Gómez et al., 2019). This store underlies transient alterations in cytosolic and lumenal Ca^2+^ that appears to contradict the principle of mass conservation. The physical basis of this store is thought to be Ca^2+^ stored via oligomerized CASQ1 (Guerrero-Hernandez et al., 2010; Perez-Rosas et al., 2015; Sánchez-Gómez et al., 2019). LLPS is an ideal mechanism to explain this behavior as a great deal of Ca^2+^ bound within the LLPS body cannot freely exchange with the surrounding aqueous environment. We posit only the periphery of the LLPS body (termed the “Diffusional Layer”, Figure 4H) can freely exchange. Ca^2+^ bound at the center of the LLPS body is necessarily less capable of exchanging with the aqueous milieu and, given its phase separation, is effectively sequestered and concealed (Figure 4H). LLPS bodies can be localized to and accessed by ryanodine receptors without dramatic alterations in local free Ca^2+^ concentrations given the reduced Ca^2+^ exchange. LLPS would allow energy intensive Ca^2+^ import machinery, like SERCA, to rapidly work against less steep Ca^2+^ gradients than total lumenal Ca^2+^ concentrations and oligomerized but solvent exposed CASQ1 would allow. LLPS, therefore, represents both a mechanism to store large quantities of Ca^2+^ but also to hold lumenal free Ca^2+^ at constant levels in a kinetically and energetically efficient manner.

FAM20C is an important regulator of Ca^2+^ handling both *in vivo* and in cell culture (Pollak et al., 2018; Worby et al., 2021). Data presented here suggest that this function is exerted, at least in part, through modulation of the intrinsic disorder and LLPS activity of CASQ1 as it is expressed in various tissues and central to excitation-contraction coupling in skeletal muscle (Woo et al., 2020). FAM20C may similarly regulate other ER/SR Ca^2+^ storage proteins, many of which are also characterized by low iso-electric point and intrinsic disorder. Confirmed ER/SR Ca^2+^ storage protein substrates include Calreticulin (CALR) and cardiac specific Calsequestrin 2 (CASQ2) (Pollak et al., 2018). Parallel to its action against ER/SR resident proteins, FAM20C phosphorylates a number of intrinsically disordered secreted proteins known to bind Ca^2+^ or calcium salts and undergo calcium-dependent association (Tagliabracci et al., 2012, 2015). These include caseins and the SIBLING proteins including Osteopontin. Both casein and Osteopontin bind Ca^2+^/calcium salts and prevent formation of biologically inaccessible and toxic calcium precipitates. This is important to neonatal nutrition and biomineralization and disruption of FAM20C activity has been linked to various disease states (Grzybowska, 2018; Kalmar et al., 2012). Indeed, loss of FAM20C activity results in a rare genetic condition termed Raine syndrome characterized by widespread ectopic calcification, which we hypothesize is directly linked to Ca^2+^/calcium salt misregulation through the disruption of LLPS (Eltan et al., 2020). The ability of FAM20C to regulate intrinsic disorder and the LLPS of these proteins is a plausible mechanism to explain these phenotypes as LLPS would allow these proteins to reversibly bind Ca^2+^ and increase local Ca^2+^ concentrations while simultaneously preventing calcium precipitation.

We hypothesize Ca^2+^-dependent LLPS is a widespread and evolutionarily conserved phenomena. We show for the first time that Ca^2+^-dependent LLPS is reversible and occurs within cells, however, liquid-like states are observed for other calcium-protein interactions *in vitro*. Polymer-Induced Liquid Precursors (PILP) occur when protein interacts with emulsions of calcium salts to enter a transient liquid-like state in the process of mesocrystallization (Wolf et al., 2011). In biomineralization, mesocrystallization is regulated by SIBLING proteins including Osteopontin (Wojtas et al., 2012). PILP have been observed for proteins from various eukaryotic organisms including zebrafish and chicken (Wojtas et al., 2012; Wolf et al., 2011). A prime example of a protein displaying PILP is Starmaker, a secreted phosphoprotein involved in otolith formation in zebrafish. This protein is intrinsically disordered, hyperphosphorylated, and contains a number of FAM20C consensus phosphorylation motifs. It inhibits calcium carbonate crystallization and induces formation of amorphous calcium carbonate structures that are reminiscent of CASQ1 structures observed in EM (Figure 1A) (Wojtas et al., 2012). These parallels suggest intrinsically disordered and polyanionic proteins, including CASQ1, Starmaker, and Osteopontin, have convergent LLPS activities arising from their similar biophysical properties that are maintained across species. Similar processes may further influence the formation of well-studied Ca^2+^-dependent biological condensates like casein micelles whose exact structure, despite extensive study, remains largely mysterious (Ingham et al., 2015).

In conclusion, we identify a previously unknown mechanism of Ca^2+^-dependent LLPS occurring within the ER/SR. This activity explains discrepancies surrounding the biochemical and structural properties of CASQ1. It reveals molecular mechanisms underscoring ER/SR Ca^2+^ storage and signaling as well as complex physiological phenomena like muscle contraction. We confirm CASQ1 as a substrate for FAM20C and identify a means through which FAM20C may influence its numerous chemically and functionally similar substrates. The activity described here represents a potentially widespread Ca^2+^-dependent mechanism as supported by PILP processes in various organisms and well-studied biological condensates like the casein micelle (Ingham et al., 2015; Wojtas et al., 2012; Wolf et al., 2011). Therefore, Ca^2+^-dependent LLPS of polyanionic and intrinsically disordered proteins represents a major mechanism underlying Ca^2+^ handling and signaling.

## Methods

### Bioinformatic Search

Sequences for the human proteome were retrieved from UniProt (http://www.uniprot.org), only reviewed sequences were considered. Sequence analysis was performed in R. Isoelectric points were determined using the R package Peptides. Amino acids composition analysis was performed using the R package stringr. Gene ontology analysis was performed using the R package enrichR. Data was visualized using the R package ggplot2.

### Molecular Cloning

Human cDNAs were purchased from either Open Biosystems or DNASU. For recombinant expression in *E. coli,* CASQ1 (amino acids 35-396) was subcloned into a pet28a-derivative vector containing an N-terminal 6X Histidine tag separated from the coding sequence by a 3C protease site. For insect cell expression, FAM20C (amino acids 93-584) was subcloned into a modified pI-secSUMOstar vector (LifeSensors, Malvern, PA) in which the original SUMO tag was replaced by a MBP tag and tobacco etch virus (TEV) protease site as previously described (Xiao et al., 2013). FLAG tagged mammalian expression constructs utilized in Figure 3A were generated by subcloning CASQ1 (amino acids 1-396) into a pCDNA3 vector with a C-terminal FLAG tag as previously described (Pollak et al., 2018). HA-tagged mammalian expression constructs utilized in Figure 3A were generated by subcloning FAM20C (full length) into a pCDNA3 vector with a C-terminal HA-tag as previously described (Pollak et al., 2018). C-terminally tagged GFP mammalian cell expression constructs utilized in Figure 1B & Supplementary Figure 6C-6E were generated by subcloning CASQ1 (amino acids 1-396) into a pEGFP vector. The ssGFP-CASQ1 construct used in Figure 4G was generated by initially cloning CASQ1 (amino acids 1-396) into a pcDNA3 derivative vector lacking epitope tags. EGFP was subcloned between amino acids 49 and 50 of CASQ1. The ssGFP construct used in Supplementary Figure 6B was generated using site directed mutagenesis to remove CASQ1 (amino acids 50-396) coding regions. ssGFP-GFP used in Supplementary Figure 6B was generated by subcloning an additional EGFP sequence C-terminally to ssGFP. Site directed mutagenesis was performed to generate FAM20C D478A using QuikChange (Agilent Technologies, Santa Clara, CA) as described previously (Pollak et al., 2018). Site directed mutagenesis was performed to generate CASQ1 S248E, CASQ1 S369E, CASQ1 S248/369E, and ssGFP constructs using SLIM (Chiu et al., 2004).

### Protein Expression and Purification

His-3C-CASQ1 constructs were expressed in *E. coli* BL21 (DE3)-RILP cells. Cultures were grown at 37°C in LB media supplemented with kanamycin to an OD_600_ of 0.6*-*0.8 and expression was induced by addition of isopropyl-β-D-1-thiogalactopyranoside (IPTG) to 400 μM. Induced cultures were grown at 25°C overnight and cells were harvested by centrifugation at 4,680xg for 15 minutes. Cell pellets were resuspended in lysis buffer (50mM Tris-HCl pH 8.0, 500mM NaCl, 15mM imidazole, 10% glycerol, 0.1% Triton X-100, and 10mM β-mercaptoethanol) and lysed via sonication. The lysate was cleared by centrifugation at 11,950xg for 45 minutes. The cleared lysate was incubated with Ni-NTA agarose (Invitrogen) for 20 minutes on ice and poured through a gravity column. The beads and bound protein were washed with wash buffer (50mM Tris-HCl pH 8.0, 500mM NaCl, 15mM imidazole, 10mM β-mercaptoethanol). Protein was eluted from the column in wash buffer containing 400mM imidazole. The solution was dialyzed against kinase reaction buffer (50mM HEPES pH 7.5, 50mM NaCl, 1mM DTT) and treated with 3C-protease at a concentration of 10μg/mL overnight at 4°C. ATP and MnCl_2_ were added to the dialyzed sample to final concentrations of 2mM. The dialyzed sample was split to generate CASQ1 and pCASQ1 reactions. Purified FAM20C was added to the pCASQ1 reaction to a final concentration of 0.007 mg/mL. The reaction was incubated at 37°C for 8 hours. Reactions were cleared to remove precipitate and CASQ1/pCASQ1 was separated from other reaction components using anion exchange chromatography. Samples were loaded onto a Resource Q column (GE Life Sciences) and fractionated using an ÄKTA FPLC (GE Life Sciences). Fractions were resolved on 15% SDS-PAGE and stained with Coomassie Brilliant Blue to confirm the presence of CASQ1/pCASQ1. Pooled anion exchange fractions were concentrated using an 30kDa MWCO Amicon Ultra 15 centrifugation concentrator (Millipore) to ∼1mL. Concentrated sample as loaded onto a HiLoad 16/600 Superdex 200 pg column (Cytiva) and fractionated using an ÄKTA FPLC (GE Life Sciences). Fractions were resolved on 15% SDS-PAGE and stained with Coomassie Brilliant Blue to confirm the presence of CASQ1/pCASQ1. Fraction were pooled and concentrated as before to ∼10mg/mL as determined by Nanodrop (Thermo) absorbance at 280nm with correction by calculated extinction coefficient from amino acid sequence. Homogeneity of phosphorylated protein was confirmed with SDS-PAGE gel shift, melting temperature determination, and analytical size exclusion chromatography.

MBP-tagged Fam20C protein was expressed in Hi-5 insect cells and was purified as described previously (Xiao et al., 2013). The MBP tag was removed via gel filtration chromatography following TEV protease cleavage.

### Electron Microscopy

CASQ1 samples for Electron Microscopy analysis were oligomerized in a solution containing 4μM CASQ1, 10% glycerol, 100mM KCl, 10mM Tris-HCl pH 7.0, and 10mM CaCl_2_ and incubated at room temperature for 10 minutes and transferred to ice. 20μL of the oligomerized sample was added dropwise to a 100 mesh formvar grid and incubated for 10 minutes to allow for adherence. The grid was washed by incubating grid in a water droplet for 1 minute. This was repeated twice for 2 minutes in fresh water droplets. Transferred grid to a droplet of 2% uranyl acetate and incubate with gentle agitation for 1 minute. Excess stain was blotted away and the mesh was allowed to dry for 5 minutes. Electron micrographs were gathered on a JEOL 1200 EX II TEM using standard settings (80kV and 40μA). Experiments were performed in collaboration with Electron Microscopy Facility at UCSD.

### Mammalian Cell Culture and Transfection

HEK293T, U2OS, and HEK293A cells were cultured in Dulbecco’s modified Eagle’s medium (DMEM, Life Technology) supplemented with 10% FBS (Life Technology) and 100μg/mL penicillin/streptomycin (Life Technology) at 37°C in 5% CO_2_. Transfections were carried out using polyethylenimine (PEI), as previously described (Longo et al., 2013).

### Live Cell Imaging

U2OS and HEK293T cells were seeded into 35mm imaging dishes at a density of 0.3×10^6^. 24 hours post seeding cells were transfected with 1μg CASQ1-GFP construct DNA. 24 hours post transfections media was replaced and cells were imaged in a Keyence BZ-9000 imaging system using an inverted 60X oil immersion lens in a channel appropriate for GFP. Thapsigargin treatment was performed by direct addition of Thapsigargin to cells to a final concentration of 5μM. Cells were observed over a 10-30 minute time frame until the majority of puncta were dissolved. Images were taken every 0.5-1 minute to confirm gradual dissolution.

### Brightfield Microscopy of LLPS CASQ1

Bright field microscopy performed in Figure 1C, 1D, and S3 were performed in a Keyence BZ-9000 imaging system using an inverted 40X or 60X oil immersion lens. Solutions imaged contained 50μM CASQ1, 10mM Tris-HCl pH 7.0, 10% glycerol, and 150mM KCl. Oligomerization was induced through addition of CaCl_2_ to 10mM. Images were taken 5-10 minutes following induction. Oligomerization was reversed through the addition of EDTA to 50mM and oligomerized species rapidly dissolved. Wetted images were taken in a focal plane allowing for visualization of the slide surface at times indicated in figure legend.

### LLPS Phase Diagram Generation

Solutions of CASQ1/pCASQ1 and calcium were combined to yield final concentrations indicated. Reactions were performed in a 2μL volume on the imaging surface of a 35mm imaging dish. Reactions contained indicated concentrations of CASQ1/pCASQ1 and calcium as well as 150mM KCl, 10mM Tris-pH 7.0, and 10% glycerol. Reactions were allowed to oligomerize for 5 minutes and then imaged in a Keyence BZ-9000 imaging system using an inverted 60X oil immersion lens. Samples were visually inspected exampled for liquid-liquid phase separated droplets. Reactions were paired in time and temperature.

### Analytical Size Exclusion Chromatography

A Superdex 200 10/300 GL (Cytiva) column was equilibrated with buffer containing 10mM Tris-HCl pH 7.0, 10% glycerol, and indicated concentration of KCl. For experiments including trifluoperazine (TFP) (Figure 3N & 3O) the column was equilibrated with 10mM Tris-HCl pH 7.0, 10% glycerol, 150mM KCl, and 100μM TFP or vehicle (DMSO). Samples were loaded onto the column and separated at a flow rate of 0.5mL/minute on an ÄKTA FPLC (GE Life Sciences). Elution was monitored by absorbance at 280nm. Stokes radii was determined from a standard curve of Gel Filtration Standards (BioRad) with known Stokes radii. Average elution volume as determined from maximal peak in absorbance at 280nm trace. All experiments were conducted at 4°C.

### Tryptophan Fluorescence Spectroscopy

Equivalent sample volumes containing 1μM CASQ1/pCASQ1, 10mM Tris-HCl pH 7.0, and indicated KCl concentration were monitored in an infinite M200PRO plate reader (Tecan). Gain and Z-distance were optimized across all samples. Samples were excited at 295 nm and emission was observed from 320 to 400nm with a 2nm step and 20nm bandwidth. Readings were performed at ambient temperature.

### Limited Proteolysis

Limited was performed in reactions containing 10mM Tris-HCl pH 7.0, 10% Glycerol, 0.3mg/mL CASQ1 or pCASQ1, 3μg/mL α-chymotrypsin (Sigma), and the indicated concentration of KCl at 30°C. Reactions were initiated by mixing a 2X solution of CASQ1 or pCASQ1 with a 2X solution of α-chymotrypsin. All reagents were preincubated to 30°C. Fractions were collected at indicated time points and mixed with an equal volume of 2X SDS-PAGE loading dye. Samples were immediately boiled at 95°C to terminate the reaction. Samples were allowed to cool and a volume corresponding to 3μg of CASQ1 or pCASQ1 was resolved on a 15% SDS-PAGE gel. Gels were stained with Coomassie Brilliant Blue and imaged using a ChemiDoc (BioRad).

### Turbidity Assay

Turbidity assays were carried out in solutions containing 4μM CASQ1/pCASQ1/CASQ1 mutant, 10mM Tris-HCl pH 7.0, 10% glycerol, indicated concentrations of CaCl_2_, and indicated concentration of KCl. Reactions were initiated by mixing a 2X solution of protein with a 2X solution of CaCl_2_. Reactions were allowed to oligomerize for 10 minutes and absorbance at 350nm readings of equivalent volumes were taken in a Spark plate reader (Tecan) using default Absorbance settings. All reactions were carried out at ambient temperature.^32^

### P Orthophosphate Metabolic Labelling

HEK293 cells were seeded at 5×10^5^ cells per well in a 6-well plate; 20 hr later, cells were transfected with indicated plasmids. 1 day after transfection, metabolic labeling was started by replacing the medium with phosphate-free DMEM containing 10% dialyzed FBS and 1mCi/mL ^32^P orthophosphate (PerkinElmer, Waltham, MA). After labeling for 8 hours, the cell lysate was collected and the cell debris was removed by centrifugation. FLAG-tagged proteins were immunoprecipitated from the supernatant and analyzed for protein and ^32^P incorporation by immunoblotting and autoradiography.

### *In vitro* Phosphorylation Assay

Proteins and indicated kinase were incubated in 50mM Tris-HCl pH 7.0, 10mM MnCl_2_, 1mM [γ-^32^P] ATP (specifi activity = 100–500cpm/pmol), 0.25mg/mL susbtrate, and 10μg/mL FAM20C at 30°C as previously described (Tagliabracci et al., 2012). Reactions were terminated at 20 minutes by SDS loading buffer, 15mM EDTA, and boiling. Reaction products were separated by SDS-PAGE and incorporated radioactivity was visualized via autoradiography and immunoblotting.

### Differential Scanning Fluorimetry

Differential scanning fluorimetry was performed in a C1000 Touch Thermal Cycler CFX96 Real-Time System (BioRad). 20μL reactions contain 1μg CASQ1/pCASQ1/CASQ1 mutant, 1X SYPRO Orange (diluted from 5000X stock in DMSO, Sigma), 10mM Tris-HCl pH 7.0, 500mM KCl (to limit oligomerization), and indicated calcium concentration (if relevant). Reactions were heated from 4°C to 95°C with fluorescence monitored every 0.5°C in both the HEX and FAM channels. Melting temperature was determined from the region of maximal slope as visualized by the first derivative of fluorescence with respect to temperature (minimal -d(RFU)/dT) per default behavior of Bio-Rad CFX manager software (BioRad).

### Sedimentation Assay

Oligomerization assays were performed as described above with slight modifications. Reactions were performed at 30°C and at a volume of 100μL. Samples were allowed to oligomerize for 20 minutes and the dense phase was separated from the light phase by centrifugation at 13,000xg for 5 minutes. The supernatant light phase was carefully aspirated and the volume was noted. The pelleted dense phase was resuspended in an equivalent volume of buffer. 0.5M EDTA, pH 8.0 was added to each sample to a final concentration of 50mM and 5X SDS-PAGE loading dye was added to a final 1X concentration. Samples were boiled for 5 minutes at 95°C. 10μL fractions of the cooled samples were resolved on 15% SDS-PAGE and stained with Coomassie Brilliant Blue stain. Protein concentration was determined from quantification of band intensity using ImageJ and converted to percentage of total protein.

### Protein Crystallization, Data Collection, and Refinement

CASQ1 S248E crystals were grown using vapor diffusion method in sitting-drop at room temperature by mixing 1μL of ∼ 10mg/mL protein with 1μL of mother liquor. Optimal CASQ1 S248E crystals were obtained using a mother liquor containing 0.1M sodium acetate, 8% PEG 400, pH 4.1. The crystals were cryoprotected in mother liquor containing 30% glycerol, flash frozen and shipped to Advanced Light Source (ALS) for data collection. Data were collected on Beamline 821. Diffraction images were processed using iMosflm in CCP4. CASQ1 S248E structures were determined by molecular replacement using previously solved CASQ1 structures (PDB ID: 5CRD) in Phenix Phaser-MR. The molecular replacement solutions were iteratively improved using Coot and Phenix Refine. The quality of the final refined structure was evaluated using MolProbity. Structures were visualized using PyMol. The final statistics for data collection and structure determination are summarized in Supplementary Table 1.

### Homo-FRET Assay

HEK293A cells were transfected with indicated plasmids. 24h post transfection cells were rinsed with 1X DPBS, harvested with mechanical scraping, and resuspended as individual cells in 1X Hanks’ Balanced Salt Solution (Millipore SIGMA). Cell concentration was determined using trypan blue staining in a Countess II cell counter (Life Technologies). 1×10^4^ to 1×10^5^ cells were loaded into the wells of a black 384-well plate, consistent cell counts were used for compared experiments. Fluorescence polarization readings were performed in a Spark plate reader (Tecan) according to default settings. Samples were excited with at 485nm (20nm bandwidth) and emission was detected at 535nm (20nm bandwidth). Optimal gain was determined automatically. Samples were excited with 200 flashes. Lag time was set to 40μs. Experiments were performed at 25 °C. For cells treated with thapsigargin and/or calcium chloride cells were automatically injected with HBSS containing sufficient amounts of thapsigargin and calcium to bring final concentrations to 5μM and 1mM, respectively. For lysis experiments sodium dodecyl sulfate (SDS) was added to 1% and EDTA was added to 10mM and cells were incubated with gentle shaking for 30 minutes.

### Statistical Analysis and Data Visualization

Unless otherwise noted all data analysis was performed in R and data was visualized using the R package ggplot2.

## Acknowledgments

The authors thank Jack Dixon for mentorship, support, and scientific guidance through the lifetime of this project. We would like to thank Jian Wu and Susan Taylor for assistance with crystal shipping and coordination with Advanced Light Source. We would like to thank Timo Meerloo and Ying Jones of the UCSD Electron Microscopy Facility for help with data collection and sample preparation.

## Author contributions

Conceptualization: JEM

Methodology: JEM

Investigation: JEM, AJP, CAW, JCX, VT

Visualization: JEM

Funding Acquisition: JEM, CAW, VT, ACN, JEM

Project Administration: JEM

Supervision: ACN, JEM

Writing – original draft: JEM

Writing – review & editing: JEM, AJP, CAW, JCX, VT, CAN

## Data and material availability

X-ray crystallography data is available in the Protein Data Bank (PDB) under accession code 8F48.

