## Supplementary materials for "Ca^2+^-dependent liquid-liquid phase separation underlies intracellular Ca^2+^ stores"

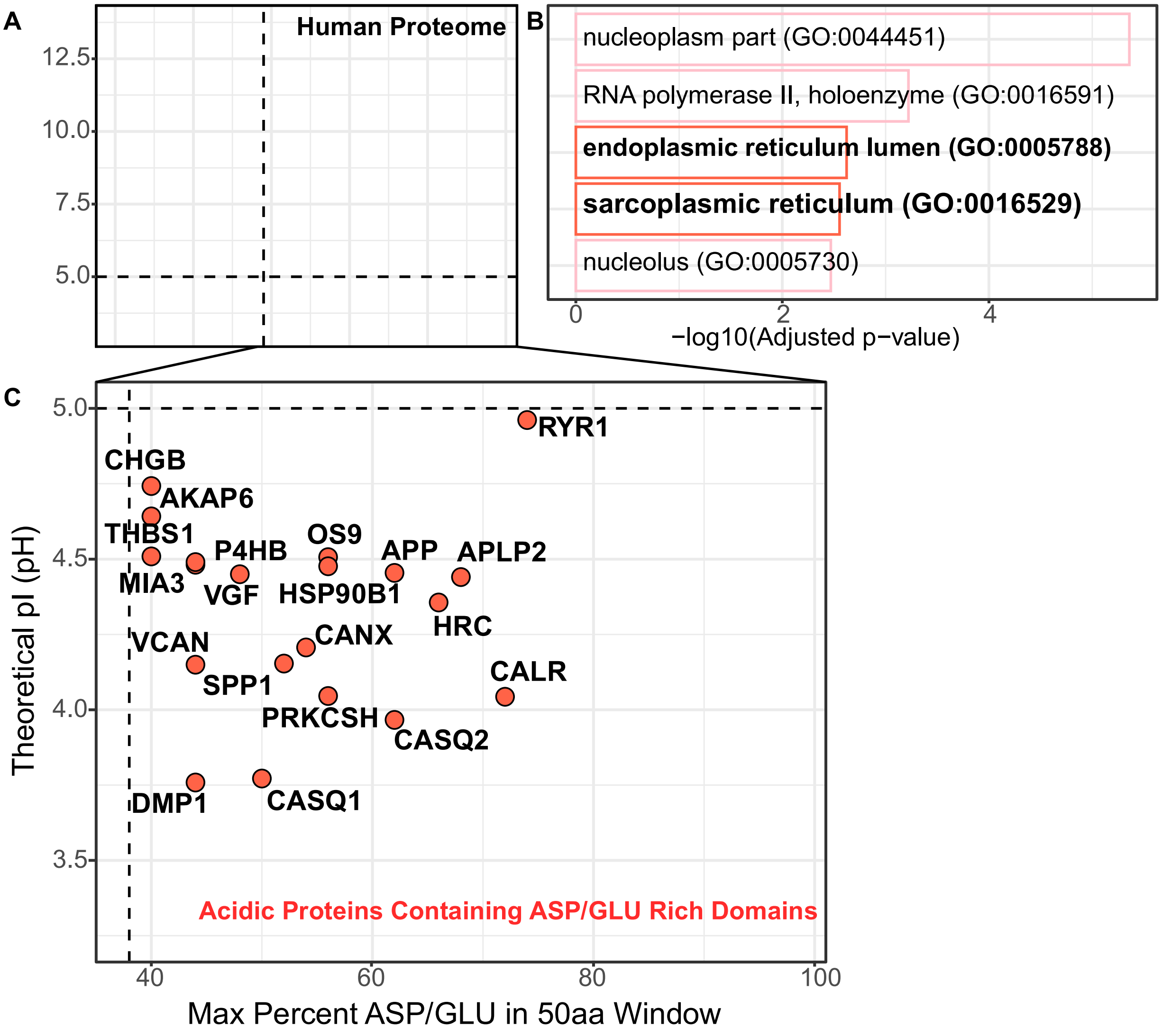


**Supplementary Figure 1.**

**Human proteome-wide sequence analysis. (A)** Proteins stratified by predicted isoelectric point and max percent acidic residues in a 50 amino acid window along the amino acid sequence. Red points indicate protein meeting criteria (>40% aspartate/glutamate in any 50 amino acid window and an overall isoelectric point below 5). Sequences retrieved from UniProt (https://www.uniprot.org). **(B)** EnrichR GO Cellular compartment ontology analysis reveals acidic proteins with highly acidic regions localize primarily to the nucleus and ER/SR. **(C)** Proteins with acidic regions and acidic isoelectric points. Labelled large red points indicate either FAM20C substrates or proteins with functional relevance to ER/SR calcium handling.

A

B

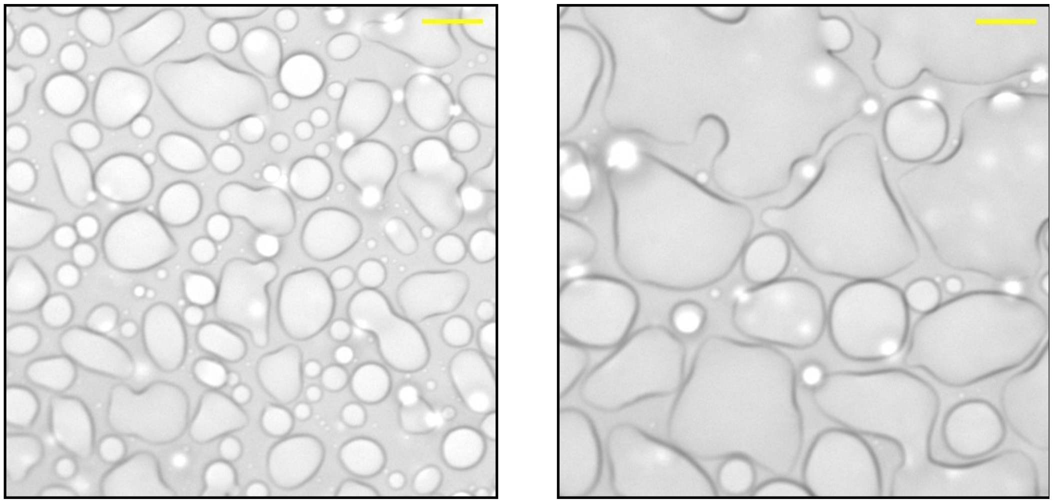

**Supplementary Figure 2.**

**CASQ1 Wetting**. **(A)** CASQ1 LLPS wetting of slides in experiments similar to those performed in Figure 1C & 1D. Image taken approximately 20 minutes following induction of oligomerization. (scale bar (yellow) = 10 μm) **(B)** Time progressed image demonstrating the gradual growth of wetted bodies. Image taken approximately 40 minutes following induction of oligomerization. (scale bar (yellow) = 10 μm).

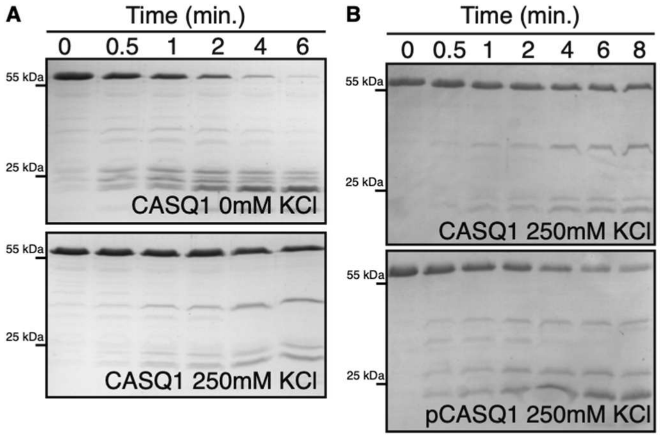


**Supplementary Figure 3.**

**Limited Proteolysis of CASQ1. (A)** Limited proteolysis of CASQ1 by chymotrypsin at different ionic strengths: 0mM KCl (top) and 250mM KCl (bottom). Time points indicated. Fractions removed from continuous reaction. **(B)** Limited proteolysis of CASQ1 by chymotrypsin in the unphosphorylated (CASQ1, top) and phosphorylated (pCASQ1, bottom) states. Ionic strength held constant at 250mM. Fractions removed from a continuous reaction.

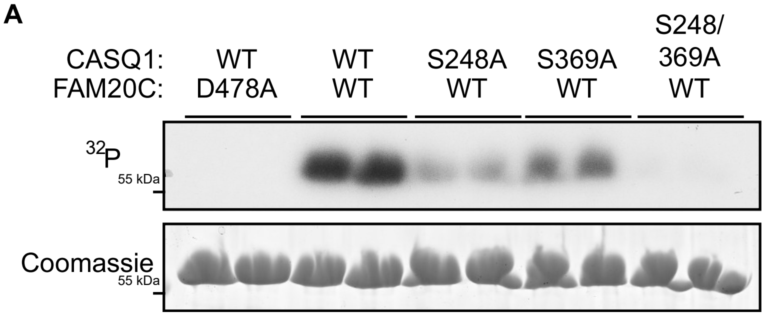


**Supplementary Figure 4.**

***In vitro* phosphorylation of CASQ1 with FAM20C. (A)** CASQ1 WT and CASQ1 alanine mutants (S248A, S369A, and S248/369A - indicated) were incubated with WT or catalytically inactive (D478A) FAM20C (indicated) in the presence of [γ-$^{32}$P] ATP. Fractions of each reaction resolved on SDS-PAGE. Representative autoradiograph (top) and Coomassie staining of SDS-PAGE gel (bottom). Two independent reactions were loaded for each substrate.

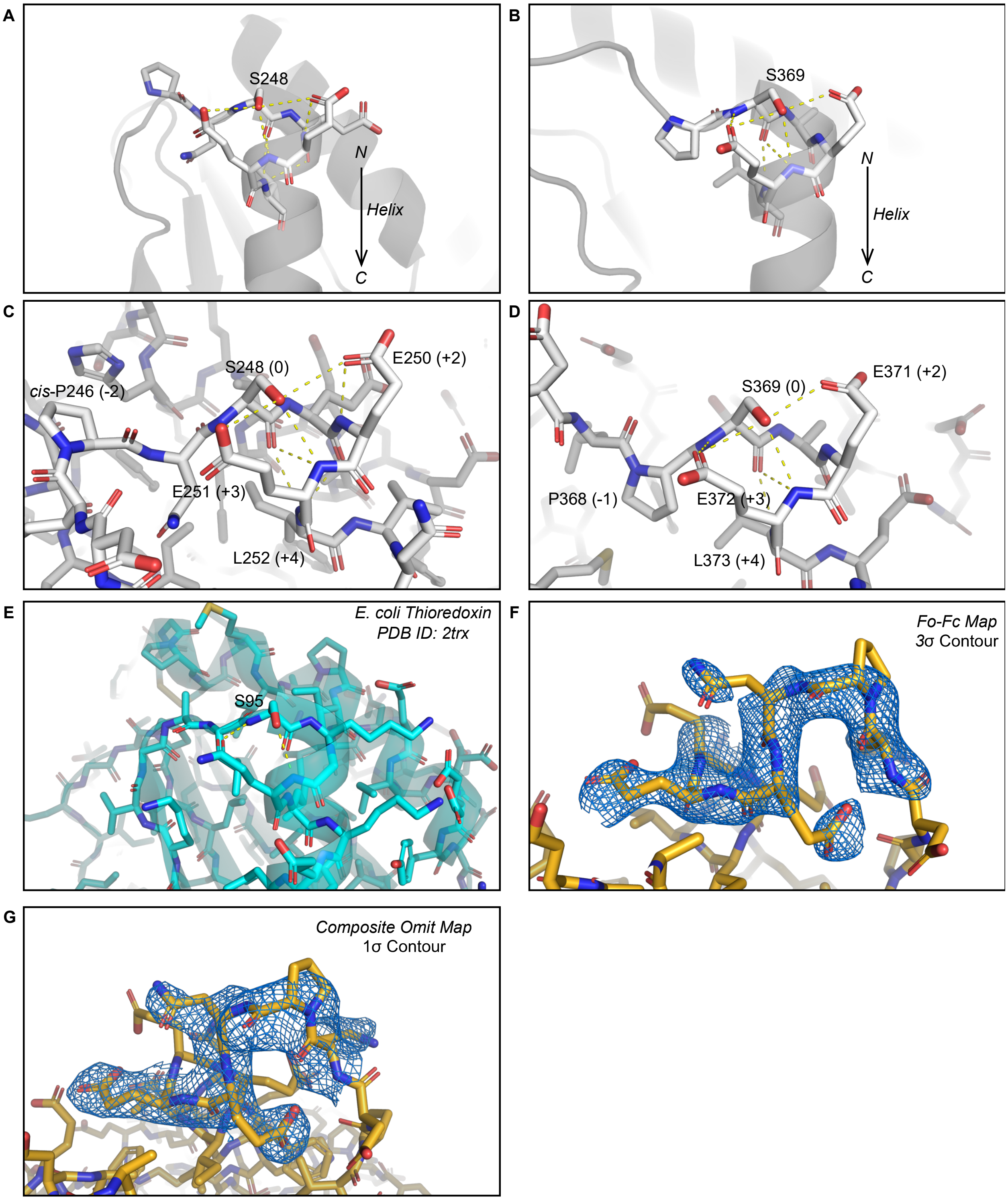


**Supplementary Figure 5.**

**Structural analysis of CASQ1 and CASQ1 S248E. (A)** Helix cap formed by S248 in CASQ1 WT (PDB ID: 5CRD). Yellow dashed lines indicate polar contacts. Arrow indicates directionality of the helix N-terminus to C-terminus. **(B)** Helix cap formed by S369 in CASQ1 WT (PDB ID: 5CRD). Yellow dashed liens indicate polar contacts. Arrow indicates directionality of the helix N-terminus to C-terminus. **(C)** Detailed view of helix cap formed by S248 in CASQ1 WT (PDB ID: 5CRD). Yellow dashed lines indicate polar contacts. Important residues and their relative distance from S248 indicated in parenthesis. **(D)** Detailed view of helix cap formed by S369 in CASQ1 WT (PDB ID: 5CRD). Yellow dashed lines indicate polar contacts. Important residues and their relative distance from S369 indicated in parenthesis. **(E)** Structurally equivalent region of *E. coli* thioredoxin domain. Polar contacts indicated by yellow dashed line. (PDB ID: 2TRX). **(F)** Fo-Fc map calculated for S248E mutation and surrounding residues. Contoured to 3σ. **(G)** Composite omit map calculated for S248E mutation and surrounding residues. Contoured to 1σ.

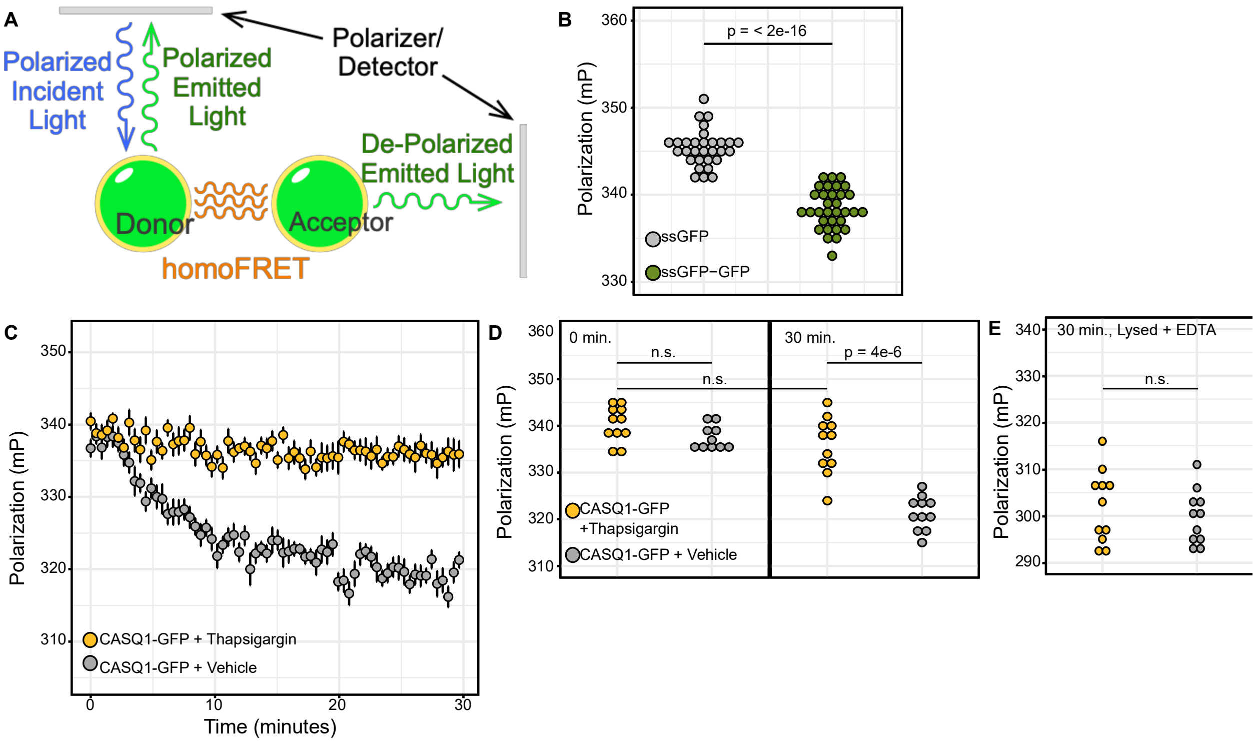


### Supplementary Figure 6.

**Homo-FRET verification in HEK293A cells. (A)** Diagram for homo-FRET. **(B)** Homo-FRET analysis of ssGFP and ssGFP-GFP. Statistical significance determined using Welch's t-test (n(ssGFP)=30, n(ssGFP-GFP)=32, alpha = 0.05). **(C)** Time-course homo-FRET measurements of CASQ1-GFP oligomerization following calcium depletion and reintroduction with (yellow) or without (grey) thapsigargin (n=11). Large points indicate time-point averages, smaller dots indicate individual data points. Error bars represent standard error of the mean. **(D)** Analysis of 0 and 30 minute time points from experiments performed in Figure 6C. Statistical significance determined using Welch's t-test (n=11, alpha = 0.05, n.s. = not significant). **(E)** Homo-FRET analysis of CASQ1-GFP following lysis and calcium chelation for cells monitored in Figure 6C-D. Statistical significance determined using Welch's t-test (n=11, alpha = 0.05, n.s. = not significant).

| | CASQ1 S248E |
|---|---|
| **PDB Code** | 7MS4 (update: 8F48) |
| **Data Collection** | |
| Wavelength (Å) | 0.97648 |
| Resolution range (Å) | 48.08 - 2.9 (3.004 - 2.9)* |
| Space group | P 21 21 21 |
| Unit cell (Å, °) | 63.65 107.34 108.17 90 90 90 |
| Total reflections | 33640 (3213) |
| Unique reflections | 16929 (1637) |
| Multiplicity | 2.0 (2.0) |
| Completeness (%) | 99.50 (98.44) |
| Mean I/sigma(I) | 9.39 (3.18) |
| Wilson B-factor | 51.28 |
| R-merge | 0.03608 (0.1111) |
| R-meas | 0.05103 (0.1572) |
| R-pim | 0.03608 (0.1111) |
| CC1/2 | 0.996 (0.962) |
| CC* | 0.999 (0.99) |
| **Refinement** | |
| Reflections used in refinement | 16928 (1637) |
| Reflections used for R-free | 1678 (157) |
| R-work | 0.1869 (0.2396) |
| R-free | 0.2966 (0.3684) |
| CC(work) | 0.946 (0.899) |
| CC(free) | 0.883 (0.726) |
| Number of non-hydrogen atoms | 5585 |
| macromolecules | 5542 |
| solvent | 43 |
| Protein residues | 680 |
| RMS(bonds) | 0.009 |
| RMS(angles) | 1.12 |
| Ramachandran favored (%) | 93.9 |
| Ramachandran allowed (%) | 6.1 |
| Ramachandran outliers (%) | 0 |
| Rotamer outliers (%) | 0.33 |
| Clashscore | 10.51 |
| Average B-factor | 46.82 |
| macromolecules | 46.91 |
| solvent | 35.23 |

*Highest resolution shell is shown in parentheses.

**Supplementary Table 1. Data collection and refinement statistics.**
